# Maternal High-Fat Diet Induces Sex- and Estrous Cycle-Specific Glial Dysregulation in the Peripheral Offspring Retina

**DOI:** 10.64898/2026.02.06.704048

**Authors:** Gintare Urbonaite, Patricija Cepauskyte, Neda I. Biliute, Guoda Laurinaviciute, Urte Neniskyte

## Abstract

**Purpose:** Maternal high-fat diet (mHFD) induces metabolic disturbances that lead to inflammatory responses in the offspring’s brain. The retina, as part of the central nervous system, may be similarly affected. This study aimed to determine how mHFD affects microglial and Müller cell activity in the retinas of offspring and assess how these effects depend on sex and the female estrous cycle.

**Methods:** Female C57Bl/6J mice were fed a control diet (CD, 10% fat) or a high-fat diet (HFD, 60% fat) from weaning to lactation. The offspring were weaned to a normal rodent diet. Retinal structure and glial cells were assessed using immunohistochemical labeling of retinal ganglion cells, Müller glia, astrocytes, phagocytic and inflammatory markers. Observed retinal changes in female offspring were correlated with the estrous cycle stages.

**Results:** mHFD induced subtle retinal structural changes and sex-specific alterations in glial cells of offspring peripheral retina. Male offspring exhibited a reduced microglial area, accompanied by increased phagocytic capacity, whereas females showed the opposite pattern. Under mCD, the microglial area and its phagocytic and metabolic activity fluctuated with the female estrous cycle, while mHFD diminished the differences between phases. Additionally, mHFD reduced Müller glial reactivity in females, indicating disrupted glial communication.

**Conclusions:** Our findings demonstrate that mHFD has a sex-specific effect on the offspring’s peripheral retina, affecting the response of retinal microglia to female reproductive hormones.

## Introduction

A high-fat diet (HFD) is one of the primary factors contributing to the development of metabolic disorders, including obesity, glucose intolerance, insulin resistance, and chronic inflammation^1–4^. There is growing evidence that these alterations could affect not only peripheral tissues but also extend to the central nervous system^5,6^. In particular, the retina is one of the most metabolically active neural tissues, relying heavily on fatty acid stores as an energy source. Alterations in nutrient supply to the retina can drive pathological changes, suggesting that metabolic disturbances induced by HFD may also adversely affect retinal health^7^.

Although the results vary, evidence from animal studies suggests that HFD may promote retinal inflammation, oxidative stress, and neurodegenerative changes^8–10^. One of the first cells to respond to stress in the retina is the microglia. HFD can disrupt microglial function, leading to an increase in microglial soma size, a morphological shift from a ramified to an amoeboid (activated) state, and an overall increase in microglial area and cell number^11,12^. Some studies link increased microglial density and metabolic dysfunction to HFD-induced inflammation and oxidative stress in the retina, enhancing microglial phagocytosis^10,13^.

Another important type of glia that maintains retinal homeostasis and provides both metabolic and structural support to neurons is the Müller cells. This specialized columnar glia spans the entire thickness of the retina^14^. In response to retinal injury or inflammatory activation, Müller cells exhibit increased expression of glial fibrillary acidic protein (GFAP), a marker widely used to indicate gliosis^15^. There is evidence that Müller cells can signal to microglia through the extracellular release of glutamate and ATP, thereby conveying changes in neuronal activity. In return, activated microglia secrete pro-inflammatory cytokines (e.g., IL-1β, IL-6), which modulate Müller cell gliosis^15,16^. Typically, microglial activation corresponds with increased Müller cell reactivity; however, in certain pathological conditions, a reduction in Müller glial activation can be observed^17^.

Increasing evidence suggests that glial responses are sex-dependent. Animal studies have shown that microglial activation, density, and morphology, as well as Müller cell gliosis, differ between males and females, both during development and in response to inflammation^18,19^. The underlying mechanisms driving these sex-specific differences are likely linked to the influence of sex hormones. Studies suggest that testosterone in males exerts anti-inflammatory effects, attenuating microglial activation and reducing the secretion of pro-inflammatory cytokines ^20^. By contrast, in females, the inflammatory response is modulated by hormonal fluctuations across the estrous cycle and can induce changes in inflammatory processes, both in the body and in the brain^21–23^. Although estradiol predominantly exerts anti-inflammatory and neuroprotective effects, it can also enhance microglial phagocytic activity^23,24^. Microglial activity can also be modulated by progesterone, which generally has anti-inflammatory properties, suppresses microglial activation and CD68 expression, but conversely increases TSPO protein levels^25–27^.

Collectively, current knowledge indicates that HFD induces inflammation, driving a wide spectrum of structural and functional alterations in the retina. However, these studies focus on the direct effects of HFD on the individual, rather than the influence of maternal high-fat diet (mHFD) on the retina of their offspring. Meanwhile, many neurodevelopmental studies have shown that maternal diet can program the metabolic and inflammatory state of the offspring, including the sensitivity of microglia to hormonal signals. For this reason, our study aimed to evaluate the impact of mHFD on the retinal structure of offspring, as well as microglial activation and Müller glia, and whether this effect differs depending on the female estrous cycle stages.

## Methods

### Animals

C57BL/6J mice were obtained from local VU Life Science Center colonies and were kept (≥ 2 mice per cage) under controlled conditions (22 °C±1, 12 h light/dark cycle, food and water *ad libitum*). Female mice were fed either a control diet (CD), containing 10% kcal from fat (Altromin diets #C1090–10 mod), or a high-fat diet (HFD), containing 60% kcal from fat (Altromin diets #C1090–60). Mice were maintained on these diets from weaning for 10 weeks, and during the subsequent pairing, pregnancy, and lactation periods. Maternal metabolic status was evaluated by weekly body weight measurements, followed by glucose and insulin tolerance tests after 10 weeks of diet feeding, prior to pairing. Female mice were paired with homozygous male *Thy1*::EGFP (IMSR Cat# JAX:007788); *Cx3cr1*::CreER (IMSR Cat# JAX:021160); *RC*::LSL-tdTomato (IMSR Cat# JAX:007905) mice on a C57BL/6J background^28^, for the fluorescence labeling of neurons and microglia cells. Both the males before pairing and the offspring after weaning were maintained on standard rodent chow (Mucedola #NFM18). The estrous cycle stages were determined by vaginal cytology in female offspring on the day of tissue collection at 22 postnatal weeks, with crystal violet dyed smears imaged using a Nikon ECLIPSE Ti-S microscope at 20X magnification. For retinal tissue analysis, we used 5-6 male and 9-12 female offspring per each of the mCD and mHFD groups. The project was approved by the Lithuanian State Food and Veterinary Service (No. G2-139). All animal experiments adhered to the requirements of European Directive 2010/63/EU and the ARVO Statement for the Use of Animals in Ophthalmic and Vision Research.

### Retinal sample collection

Eyeballs were collected from offspring whose mothers were fed either a CD or an HFD. Animals were anesthetized intraperitoneally with a mixture of 100 mg/kg ketamine and 10 mg/kg xylazine and then transcardially perfused with 0.9% NaCl. Dissected eyeballs were fixed with 4% paraformaldehyde (PFA) for 24 hours at 4 °C. After sucrose cryoprotection, fixed eyeballs were embedded in Cryomatrix embedding medium and snap-frozen at −80 °C. Frozen samples were cryosectioned at a thickness of 15 μm using a Shandon FE & FSE cryotome.

### Immunostaining

Retinal slices were permeabilized and blocked using 20% normal goat serum (NGS) and 0.4% Triton X-100 in PBS for 2 hours at room temperature with gentle shaking. Subsequently, retinal sections were incubated overnight in 20% NGS and 0.4% Triton X-100 at 4°C with selected primary antibodies overnight followed by corresponding secondary antibodies (Table 1) for 2 hours at room temperature. Immunolabeled retina was stained with DAPI (1 μg/mL) and mounted with Mowiol medium. Retinal thickness measurement and ganglion cell counting were performed using a Leica TCS SP8 confocal microscope with a 20×/0.75 NA objective (0.568 μm/pixel). Anti-Iba1-marked microglia and anti-GFAP-marked Müller cells were imaged using an EVOS FL auto cell imaging system with a 20× objective (0.45 μm/pixel). Anti-tdTomato, anti-CD68, and anti-TSPO immunolabeled areas were imaged using an Olympus IX83 widefield fluorescence microscope with a 20× objective (0.325 μm/pixel).

**Table 1.** List of antibodies used in this study.

| Type | Antibody | Source / Catalog No. | RRID | Dilution |
| --- | --- | --- | --- | --- |
| Primary antibody | Rabbit anti-Iba1 | FUJIFILM Wako Chemicals U.S.A., Cat# 019-19741 | AB_839504 | 1:1000 |
|  | Mouse anti-RFP (recognizing tdTomato) | ROCKLAND, Cat# 200-301-379 | AB_2611063 | 1:1000 |
|  | Rat anti-CD68 | Thermo Fisher Scientific, Cat# 14-0681-82 | AB_2572857 | 1:200 |
|  | Rabbit anti-PBR/TSPO | Abcam, Cat# ab109497 | AB_10862345 | 1:400 |
|  | Chicken anti-GFAP | Abcam, Cat# ab4674 | AB_304558 | 1:1000 |
|  | Rabbit anti-RBPMS | GeneTex, Cat# GTX118619 | AB_10720427 | 1:1000 |
| Secondary antibody | Goat anti-rabbit Alexa Fluor 594 | Thermo Fisher Scientific, Cat# A-11012 | AB_2534079 | 1:500 |
|  | Goat anti-mouse Alexa Fluor 647 | Thermo Fisher Scientific, Cat# A-32733 | AB_2535813 | 1:500 |

### Retinal layer thickness and ganglion cell count assessment

To separate retinal layers and count ganglion cells, the slices were immunolabeled with anti-RBPMS and stained with DAPI. The thickness of retinal layers was evaluated by measuring 10-15 positions at different locations across the central and peripheral 500 µm retinal segments and calculating the average thickness for each image. Relative retinal thickness was determined according to this equation:

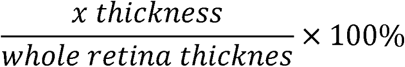

Ganglion cells labeled with anti-RBPMS were quantified by applying the same threshold and automated particle counting using ImageJ/Fiji. Cell counts were determined within a 500□ µm-long central and peripheral retinal segments (cells/500□ µm).

### Microglia and Müller cell analysis

The distribution of microglia and Müller cells in retinal segments was evaluated in Iba1 and GFAP-immunolabeled retinal slices, respectively. First, using the ImageJ/Fiji program, each 8-bit grayscale retinal image was cropped into 500 µm long segments based on the outer nuclear layer (ONL). Cell area was estimated by processing images: subtracting the background (Radius 50), applying a median filter (Radius 1), a Gaussian blur (Sigma radius 1), and finally measuring the area of the created masks using ImageJ/Fiji.

To analyze the microglia area in the central and peripheral retina, microglia cells were marked with anti-Iba1 and processed as described above. The microglia area was determined within the generated mask. Retinal sections were stained with antibodies against tdTomato (genetically labelling microglia), CD68, and TSPO and analyzed in peripheral retinal segments. Image channels corresponding to individual antibody signals were separated, and each channel was processed as described above. The same optimal threshold was selected for all retinal samples. tdTomato and CD68 or tdTomato and TSPO overlay masks were generated, and total microglial area, as well as the quantification of anti-CD68 and anti-TSPO-positive microglial areas, were performed.

Müller cell area and Müller glial processes lengths were quantified across all retinal layers in the peripheral retina. The lengths of Müller cell processes were determined at 5 different positions per retinal segment by calculating the ratio of total retinal layer width to the extent of GFAP-positive processes penetration into the layers. The channels were then split, and images were processed as described above. To determine the total GFAP-positive Müller cell area in the peripheral retina of the offspring Auto Local Threshold value was applied across all samples (Method: Bernsen; Radius: 5).

### Statistical analysis

To assess differences in area between experimental animal groups, a statistical analysis was performed using a custom Python 3.8 script generated and completed in the Spyder environment via the Anaconda platform. First, we removed outliers exceeding the mean ± 1.5 standard deviations (SD) and assessed normality using the Shapiro–Wilk test. Then, a nested two-way ANOVA was conducted to assess the effects of sex (or estrous cycle stage), maternal diet, and their interaction on the area of retinal biomarkers, including a nested random factor (Animal ID) to account for within-subject variation. Pairwise comparisons were performed using Tukey’s *post hoc* test to identify specific group differences related to sex, diet, or estrous cycle stage.

## Results

### Maternal metabolic status is altered by an HFD: effects on body mass, glucose, and insulin tolerance

In this study, to investigate how maternal diet-induced metabolic changes affect offspring retina, the mHFD mouse model was implemented (Fig. 1A). Females receiving HFD gained significantly more weight after 3 weeks of diet consumption compared to the CD group (Fig. 1B). Metabolic tests indicated that HFD females developed impaired glucose tolerance with elevated blood glucose levels from 30 min after glucose injection onwards (Fig. 1C). HFD reduced insulin sensitivity with elevated glucose levels detected at the 40- and 60-minute timepoints after insulin injection (Fig. 1D).

**Figure 1.**
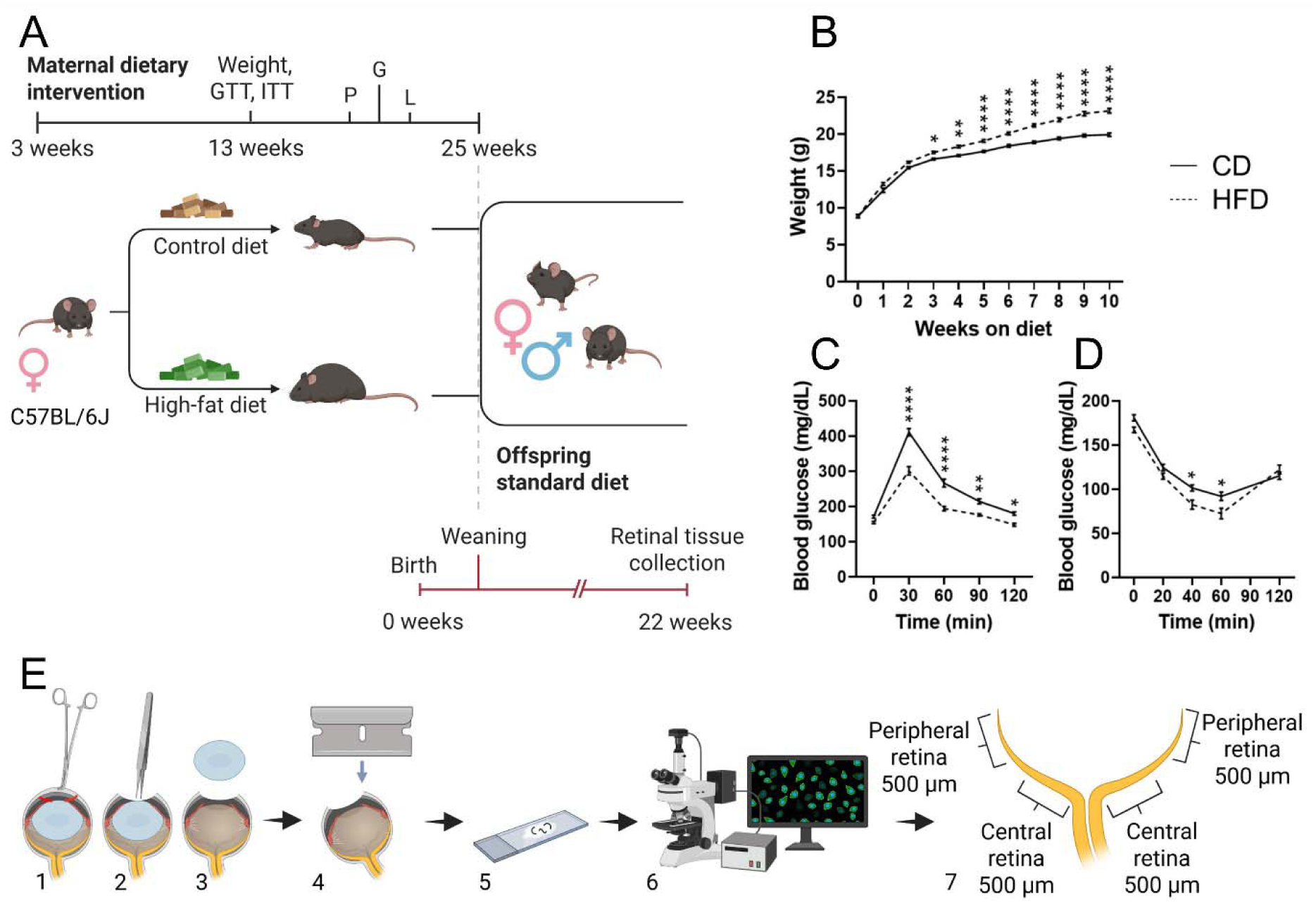
Experimental design and retinal tissue collection workflow. (A) Schematic representation of the mHFD mouse model and offspring development timeline. (B) Maternal weight. (C) Glucose tolerance test. (D) Insulin tolerance test. (E) Workflow of offspring eyeball dissection, preparation, retinal tissue collection, and fluorescence imaging. P – pregnancy, G – gestation, L – lactation.

### mHFD induced changes in peripheral retinal microglia activation depending on the sex and estrous cycle of the female

In pilot studies, microglial areas were assessed in the central and peripheral retina using Iba1 immunostaining (Supplementary Fig. S1) in samples prepared as presented in Fig. 1E. We found no differences in microglial area between groups in the central retina (Supplementary Fig. S1C); however, the effects of sex and diet were observed in the peripheral retina (Supplementary Fig. S1D). Under mCD, the microglia area in the retinas of males was larger than that of females, consistent with the known physiological effect of sex hormones^29^. However, the mHFD appears to induce a sex-specific remodeling of the microglial response, with a decrease in the male microglial area and an increase in the female microglial area (Supplementary Fig. S1D).

Based on the significant changes observed specifically in the peripheral retina, further detailed analyses were conducted in this retinal area. Using the *Cx3cr1*::tdTomato mouse, in which microglial cells are genetically labeled by tdTomato red fluorescent protein, revealed a similar significant interaction between sex and maternal diet. We found that mHFD significantly decreased the area of tdTomato+ microglia in males compared to the mCD group but increased this area in females (Fig. 2B). In this experiment, mHFD revealed a sex-specific microglial response that was not evident in the control diet group.

**Figure 2.**
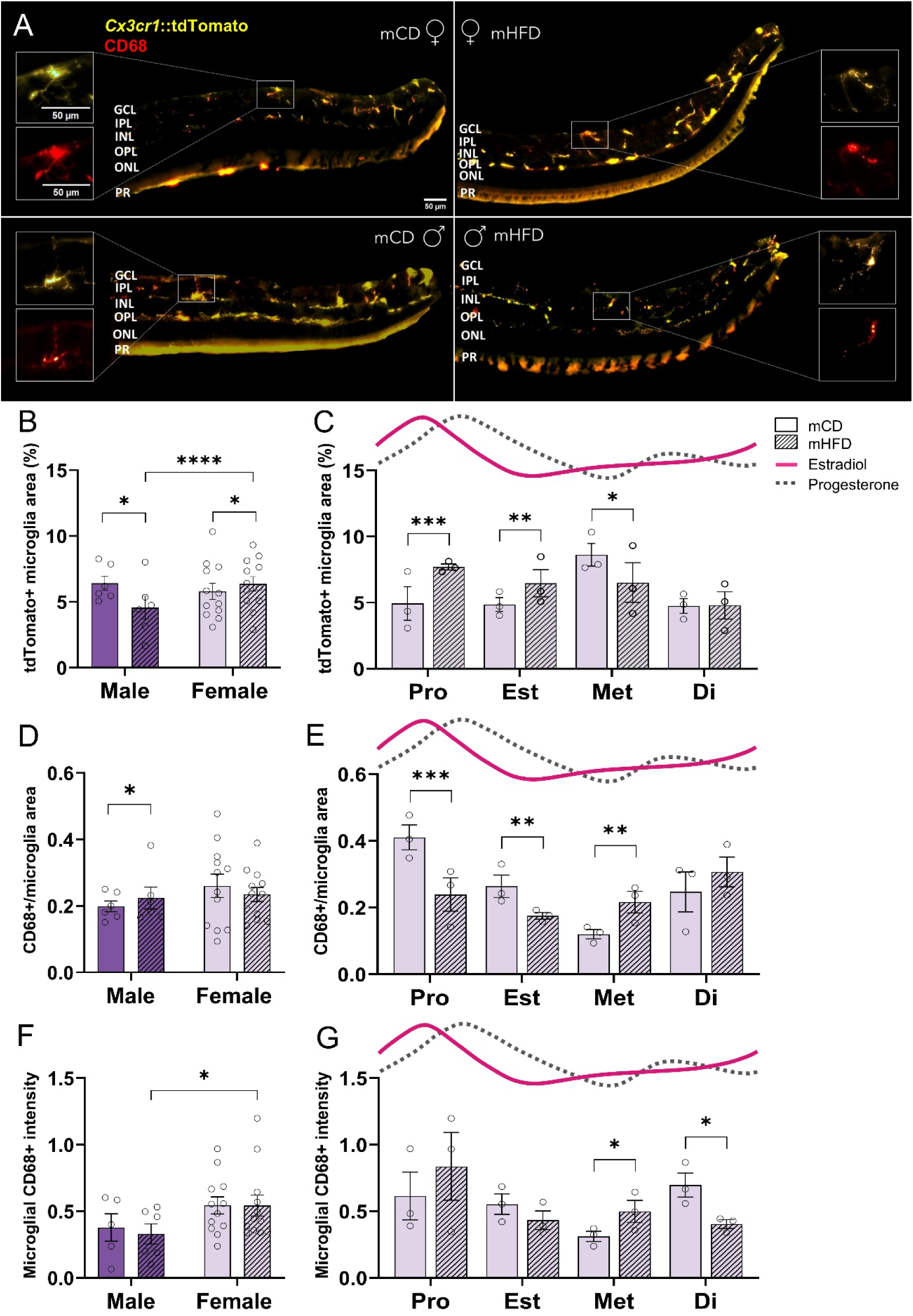
The effects of mHFD on microglia and microglial CD68 of the offspring’s peripheral retina. (A) Representative confocal images of tdTomato-positive microglia (*Cx3cr1*::tdTomato, yellow) and microglial CD68 immunoreactivity (red) in peripheral retinal sections from female (♀) and male (♂) offspring of dams who were fed either control (mCD) or high-fat diet (mHFD). GCL – ganglion cell layer. IPL – inner plexiform layer. INL – inner nuclear layer. OPL – outer plexiform layer. ONL – outer nuclear layer. PR – photoreceptor layer. Scale bar = 50 µm. (B) tdTomato+ microglia area fraction (%) by sex and (C) female estrous cycle Pro – proestrus, Est – estrus, Met – metestrus, Di – diestrus stages. Solid line – estradiol; dashed line – progesterone, based on ^30^. (D) CD68+/microglia area fraction (%) in the peripheral retina of the offspring by sex and (E) female offspring estrous cycle. (F) Offspring microglial CD68+ intensity by sex and (G) female offspring estrous cycle stages. Mean ± SEM; n = 6 males and n = 12 females per group. *P*-values were assessed using nested two-way ANOVA with Tukey *post hoc* tests. \**P* < 0.05, \*\**P* < 0.01, \*\*\**P* < 0.001, \*\*\*\**P* < 0.0001.

To address the fluctuation of female sex hormones, females were further separated by estrous cycle based on cytological images of vaginal smears (Supplementary Fig. S2). Under physiological conditions (mCD group), the area of tdTomato+ microglia varied depending on the phase of the estrous cycle – it was lower during proestrus and estrus (highest estradiol and progesterone levels), highest in the metestrus phase (lowest estradiol and progesterone levels), and then decreased again (Fig. 2C). Although hormone levels were not directly measured in this study, fluctuations in estradiol and progesterone concentrations across the estrous cycle are well documented in the literature ^30^. Importantly, we determined that mHFD disrupted these cyclical microglial dynamics, reducing the differences of microglial area between the phases (Fig. 2C). In addition, we found that mHFD significantly increased the area of tdTomato+ microglia during proestrus and estrus phases and decreased during the metestrus phase (Fig. 2C). These results suggest that mHFD alters microglia sensitivity to estrogen or progesterone signals during the estrous cycle.

One of the main functions of microglia is phagocytosis. Microglial phagocytic activity can be assessed using the cluster of differentiation 68 (CD68) glycoprotein expressed on lysosomal membranes^31,32^. Our analysis of the CD68-positive (CD68+) area of microglia revealed sex- and estrous cycle-dependent changes in microglial phagocytic capacity. As shown in Fig. 2D, we found that mHFD significantly increased the area of microglial CD68 in males, indicating enhanced microglial phagocytic activity. In females, by contrast, the CD68+ area did not increase overall, but a clear dependency on the phases of the estrous cycle was observed (Fig. 2E). After the evaluation of individual phases (Fig. 2E), we determined that the area of microglial CD68 in mCD females was the highest in the proestrus phase and the lowest during metestrus. Strikingly, we found that mHFD disrupted this cyclical pattern, reducing the area of CD68 in proestrus and estrus but increasing it in the metestrus phase (Fig. 2E). This suggests that mHFD reverses the normal direction of hormonal regulation, reducing the phagocytic activity of microglia at high estradiol and progesterone levels and increasing it when these hormones are reduced.

When we assessed the intensity of microglial CD68+ signal, we found that the total CD68 intensity was higher in females than in males, and it fluctuated during the estrus cycle in response to varying hormone levels (Figs. 2F, 2G). Meanwhile, mHFD distorted this profile – the CD68 intensity increased in metestrus and decreased in diestrus (Fig. 2G), supporting an altered microglial response to hormonal changes.

Since the changes in microglial activity may affect retinal cell viability and tissue integrity^33–35^, we assessed the thickness of the total retina and its separate layers (Supplementary Fig. S3). In mCD females, the total central retinal thickness was greater than in males, but mHFD significantly reduced it, even though this effect was not observed in males (Supplementary Fig. S3C). After the evaluation of individual central layers, we found that in males, mHFD thinned the photoreceptor layer (PR) but increased the thickness of the outer nuclear layer ONL (Supplementary Fig. S3D). In the peripheral retina, mHFD increased total retinal thickness exclusively in females (Supplementary Fig. S3E). In males, mHFD changed the thickness of individual peripheral layers, reducing the PR but increasing the thickness of the ONL and outer plexiform layer (OPL) (Supplementary Fig. S3F). Meanwhile, we did not find any effect of mHFD on the number of retinal ganglion cells in either the central or peripheral parts of the retina of either sex (Supplementary Figs. S3G, S3H). Since the effects of mHFD on microglial activity showed clear sex and hormonal cycle differences, we aimed to determine whether these changes were also reflected in other markers of microglial activation.

### mHFD altered the microglial TSPO expression in a sex- and estrous cycle-dependent manner

The maintenance of microglial function requires high levels of energy, and mitochondria play a central role in supporting these processes^36^. The translocator protein (TSPO), located in the outer mitochondrial membrane, is upregulated during retinal inflammation and serves as a marker of microglial activation^37,38^. We determined that while the microglial TSPO area in the peripheral retina was similar in both sexes of the mCD offspring, the exposure to mHFD induced significant differences in microglial TSPO between the sexes (Fig. 3B). These results are consistent with previously described differences in microglial area (Fig. 2B), indicating that both the inflammatory response of microglia and its regulation of microglial reactivity to maternal metabolic state are sex-specific.

**Figure 3.**
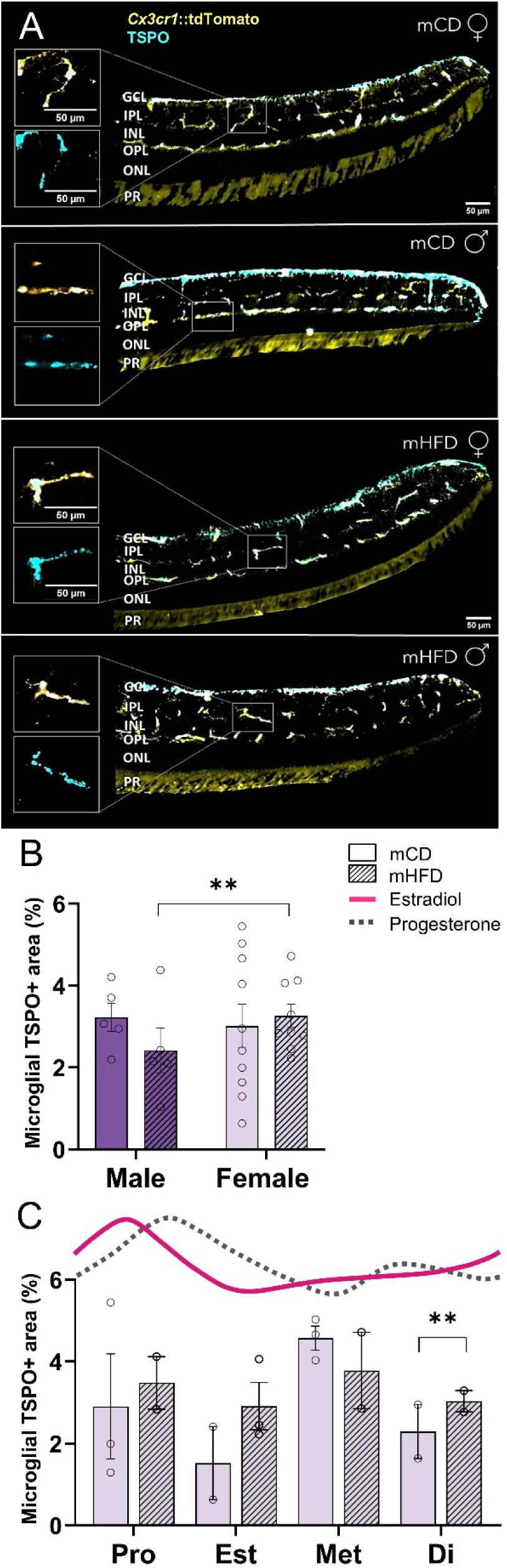
The effects of mHFD on microglial TSPO in the peripheral retina of the offspring. (A) Representative confocal images of tdTomato-positive microglia (*Cx3cr1*::tdTomato, yellow) and microglial TSPO immunoreactivity (cyan) in the peripheral retinal sections from female (♀) and male (♂) offspring of dams who were fed either control (mCD) or high-fat diet (mHFD). GCL – ganglion cell layer. IPL – inner plexiform layer. INL – inner nuclear layer. OPL – outer plexiform layer. ONL – outer nuclear layer. PR – photoreceptor layer. Scale bar = 50 µm. (B) Microglial TSPO area fraction (%) in the peripheral retina of the offspring by sex and (C) female offspring estrous cycle Pro – proestrus, Est – estrus, Met – metestrus, Di – diestrus stages. Solid line – estradiol; dashed line – progesterone, based on ^30^. Mean ± SEM; n = 5 males and n = 12 females per group. *P*-values were assessed using nested two-way ANOVA with Tukey *post hoc* tests. \**P* < 0.05, \*\**P* < 0.01, \*\*\**P* < 0.001, \*\*\*\**P* < 0.0001.

We then evaluated the retinas of females according to the phases of the estrous cycle (Fig. 3C), finding that the microglial TSPO area in the mCD group fluctuated cyclically, reflecting the changes in hormone levels. In contrast, we found that mHFD disrupted this cyclical pattern, abolishing the differences between phases and causing an increase in microglial TSPO during the diestrus phase. This suggests that mHFD distorts the hormonal regulation of TSPO expression in microglia, potentially promoting inflammatory or metabolic reactivity at phases when this activity should be physiologically suppressed. The observed changes in microglial reactivity suggest that mHFD may induce a broader glial response in the retina.

### Changes in Müller glial reactivity after mHFD exposure

To determine whether these microglial changes were accompanied by alterations in other retinal glial populations, we next assessed Müller glial reactivity. Using GFAP immunohistochemical labeling we found that mHFD tended to reduce the area of Müller glia, but a statistically significant reduction was only observed in females (Fig. 4B). Furthermore, we determined that mHFD caused shortening of the processes, also exclusively in females (Fig. 4C). These results indicate that mHFD inhibits Müller glial reactivity and may affect the structural integrity of retinal glia in a sex-specific manner. The reduction in Müller glia area and shortening of processes following mHFD suggest reduced glial reactivity. As an increase in microglial area and activity was observed simultaneously, these results suggest that mHFD induces an imbalance in retinal glial response, with microglia becoming hyperreactive and Müller glia losing their supportive and protective functions. Such an imbalance in glial interregulation may promote an inflammatory microenvironment in the retina and reduce its resistance to metabolic stress.

**Figure 4.**
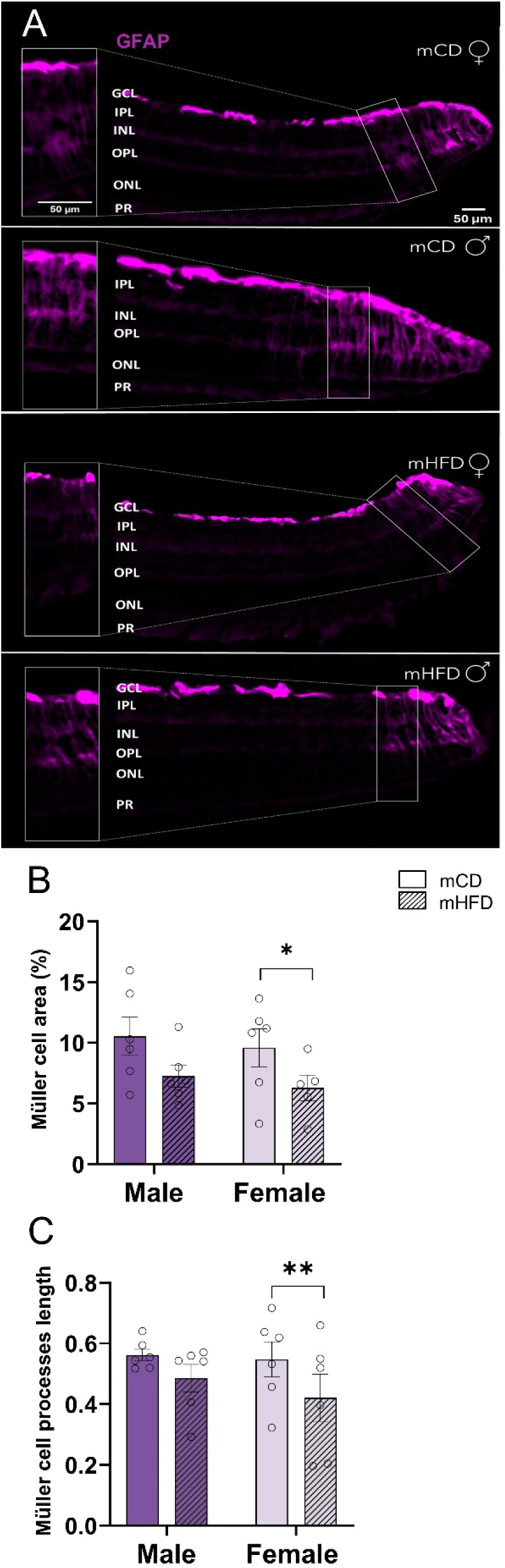
The effects of mHFD on the Müller cells in the peripheral retina of the offspring. (A) Representative confocal images of GFAP-positive (magenta) in sections of the peripheral retina from female (♀) and male (♂) offspring of dams who were fed either control (mCD) or high-fat diet (mHFD). GCL – ganglion cell layer. IPL – inner plexiform layer. INL – inner nuclear layer. OPL – outer plexiform layer. ONL – outer nuclear layer. PR – photoreceptor layer. Scale bar = 50 µm. (B) Müller cell area fraction (%) in the peripheral retina of the offspring. (C) Müller cell processes length in the peripheral retina of the offspring. Mean ± SEM; n = 5–6 per group. *P*-values were assessed using nested two-way ANOVA with Tukey *post hoc* tests. \**P* < 0.05, \*\**P* < 0.01, \*\*\**P* < 0.001, \*\*\*\**P* < 0.0001.

## Discussion

This study demonstrates that mHFD alters retinal glial function in a sex- and hormone-dependent manner, affecting both microglial activation and Müller glial reactivity. Our data are broadly consistent with previous studies showing that HFD induces inflammatory and degenerative changes in the retina. Retinal thinning, layer degeneration, and cell loss have been described in HFD models^33–35^, although some authors did not find significant differences between HFD and CD-fed mice^9,39^. In our study, we found that mHFD caused minor structural alterations in the total offspring retina and in its separate layers in both sexes (Supplementary Fig. S3). These results suggest that mHFD did not cause overt retinal degeneration in the offspring. In most studies, it was reported that HFD increased inflammation in the CNS, including in the retina, together with an increase in microglial activity^9,10^. Our results confirmed that HFD-induced inflammation is transmitted transgenerationally. We found that microglial cell area is altered in the peripheral retina (Supplementary Fig. S1), suggesting that inflammation may primarily begin in the retinal periphery. In this region, reduced vascular density and longer diffusion distances limit metabolic support, which may increase susceptibility to inflammatory activation^40^.

It is known that mHFD influences offspring brain development through inflammatory pathways, often in a sex-dependent manner. Male offspring typically show stronger pro-inflammatory microglial activation, whereas females show more variable or opposite responses^41,42^. However, sex-specific effects of mHFD on offspring retinal microglia have not been previously described. Our findings revealed that mHFD decreased the area of microglia in males but increased it in females (Fig. 2B), suggesting opposite regulation of retinal microglia between sexes that may be influenced by hormonal modulation.

Under normal conditions, we found that the area of microglia varied across the estrous cycle (Fig. 2C). This pattern aligns with studies in the hippocampus and hypothalamus, which show that high ovarian hormone levels suppress microglial activation, whereas their withdrawal promotes a more reactive morphology and increased cytokine expression^19,43^. Our results revealed that mHFD disturbed the normal pattern in the female offspring – the differences between cycle phases were reduced, and the microglial area increased when estradiol and progesterone levels were high. Together, our data suggest that maternal diet may disrupt the normal link between reproductive hormones and immune regulation in the retina, potentially increasing vulnerability to inflammation when hormonal protection is reduced. In addition to the changes in the microglia area, we observed alterations in other markers of microglial activation. Although mHFD reduced the microglial area in males, the area of phagolysosomal marker CD68 within microglia increased (Fig. 2D), indicating enhanced lysosomal or phagocytic activity. This suggests that mHFD promotes a more activated or metabolically engaged microglial phenotype even when overall microglial coverage is reduced. This pattern is consistent with findings from obesity and diabetes models, which show elevated lysosomal markers in microglia exposed to metabolic stress^13^. In mCD females, CD68 area within microglia changed dynamically across the estrous cycle (Fig. 2E), a pattern consistent with reports that ovarian hormones enhance microglial phagocytic capacity while limiting pro-inflammatory signaling^24^. However, mHFD disrupted this hormonal regulation, flattening the differences between phases. To better assess microglial activation, we also analyzed CD68 intensity (Figs. 2F, 2G), which reflects its expression level. Although the CD68 area decreased in proestrus, CD68 intensity tended to increase, suggesting a potential increase in lysosomal CD68 content per microglial cell. Together, these results suggest that mHFD alters both the extent and strength of microglial phagocytic activity, reprogramming cells toward a more reactive and hormonally unresponsive state.

In addition to the CD68 findings, we examined TSPO to further evaluate how mHFD affects microglial mitochondrial function. Under physiological conditions, microglial TSPO area followed the hormonal rhythm (Fig. 3C), showing mild fluctuations across the estrous cycle. Our results revealed that after mHFD exposure, this pattern was lost, suggesting that the normal hormonal control of mitochondrial activity in microglia was disrupted. The stronger TSPO response in females also supports a sex-dependent sensitivity of retinal microglia (Fig. 3B). It aligns with previous studies, which showed that obesity can alter microglial energy metabolism and inflammatory signaling in a sex-specific manner^44^.

Together, the changes in CD68 and TSPO indicate that mHFD promotes a more persistent microglial reactive profile that appears less responsive to hormonal regulation. Interestingly, we found that this microglial hyperreactivity was accompanied by a reduction in Müller cell area and process length in females (Figs. 4B, 4C), suggesting suppressed Müller glial reactivity. Since Müller cells play a key role in supporting neurons and communicating with microglia through ATP, glutamate, and cytokine signaling^15,16^, their reduced activation may weaken retinal homeostasis. Similar patterns, where increased microglial activity coincides with diminished GFAP expression, have been reported in neurodegenerative disease models^17,45^. Thus, mHFD appears to affect the intracellular interactions between microglia and Müller cells, but further molecular investigation is needed.

In summary, this study demonstrates that mHFD programs long-term, sex- and hormone-dependent changes in glial activity of the offspring retina. These effects primarily occur in the peripheral retina and precede overt neurodegeneration, suggesting that glial dysregulation is an early event in retinal stress. The strength of this work lies in the combined analysis of multiple glial markers (dTomato/Iba1, CD68, TSPO, GFAP), the inclusion of both sexes and estrous cycle phases, and regional evaluation across the retina, which together provide a comprehensive view of glial diversity and plasticity. However, the functional analyses, such as electrophysiology or visual behavior, were beyond the scope of this work. Future studies should address these aspects and examine whether modulation of microglial metabolism or hormone signaling can restore retinal homeostasis and reduce vulnerability to metabolic stress.

## Acknowledgements

The authors thank Igoris Nagula for the help with immunohistochemical labelling of retinal samples; Viktorija Kralikiene for technical work with animals; Dovydas Gabrielaitis, Lina Saveikyte, and Ugne Kuliesiute for their assistance with the technical setups for microscopy and image analysis.

## Author Contributions

Conceptualization: U.N.; Methodology: G.U., G.L. and U.N.; Software: G.U., P.C. and N.I.B.; Formal Analysis: G.U., P.C. and N.I.B.; Investigation: G.U., P.C. and N.I.B.; Data Curation: G.U. and U.N.; Writing – Original Draft: G.U. and P.C.; Writing – Review & Editing: G.U., P.C. and U.N.; Visualization: G.U. and P.C.; Supervision: U.N.; Funding Acquisition: G.U. and U.N.

## Declaration of Generative AI and AI-Assisted Technologies in the Writing Process

Creating Python scripts for statistical analysis, the authors used ChatGPT 4o, 5.1 versions, and Grammarly 1.142.1.0 version to improve language and grammar. After using these tools, the authors reviewed and edited the content.

