## Supplementary Fig. S1 for "Maternal High-Fat Diet Induces Sex- and Estrous Cycle-Specific Glial Dysregulation in the Peripheral Offspring Retina"

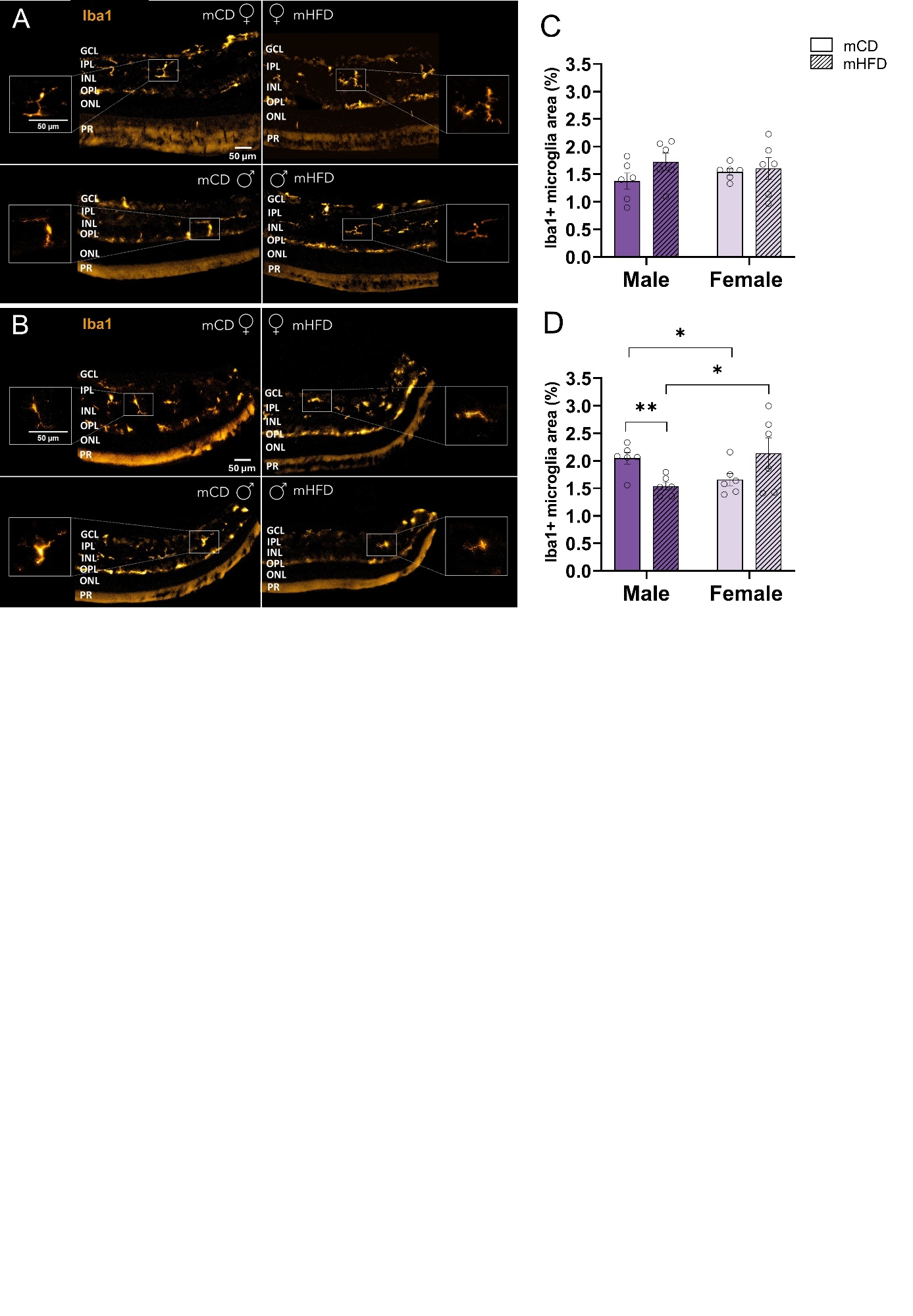

**Supplementary Figure 1.** **The effects of mHFD on the Iba1-marked microglia of the offspring retina.**  Representative confocal images of Iba1-positive microglia in central retina (A) and peripheral retina (B) sections from female (♀) and male (♂) offspring of dams who were fed either control (mCD) or high-fat diet (mHFD). (C) Iba1+ microglia area (μm²/mm²) in the central retina of the offspring. D) Iba1+ microglia area (μm²/mm²) in the peripheral retina of the offspring. Mean ± SEM, n = 6 animals per group. *P*-values were assessed using nested two-way ANOVA with Tukey *post hoc* tests. **P* < 0.05, ***P* < 0.01, ****P* < 0.001, *****P* < 0.0001.
