## Supplementary Fig. S2 for "Maternal High-Fat Diet Induces Sex- and Estrous Cycle-Specific Glial Dysregulation in the Peripheral Offspring Retina"

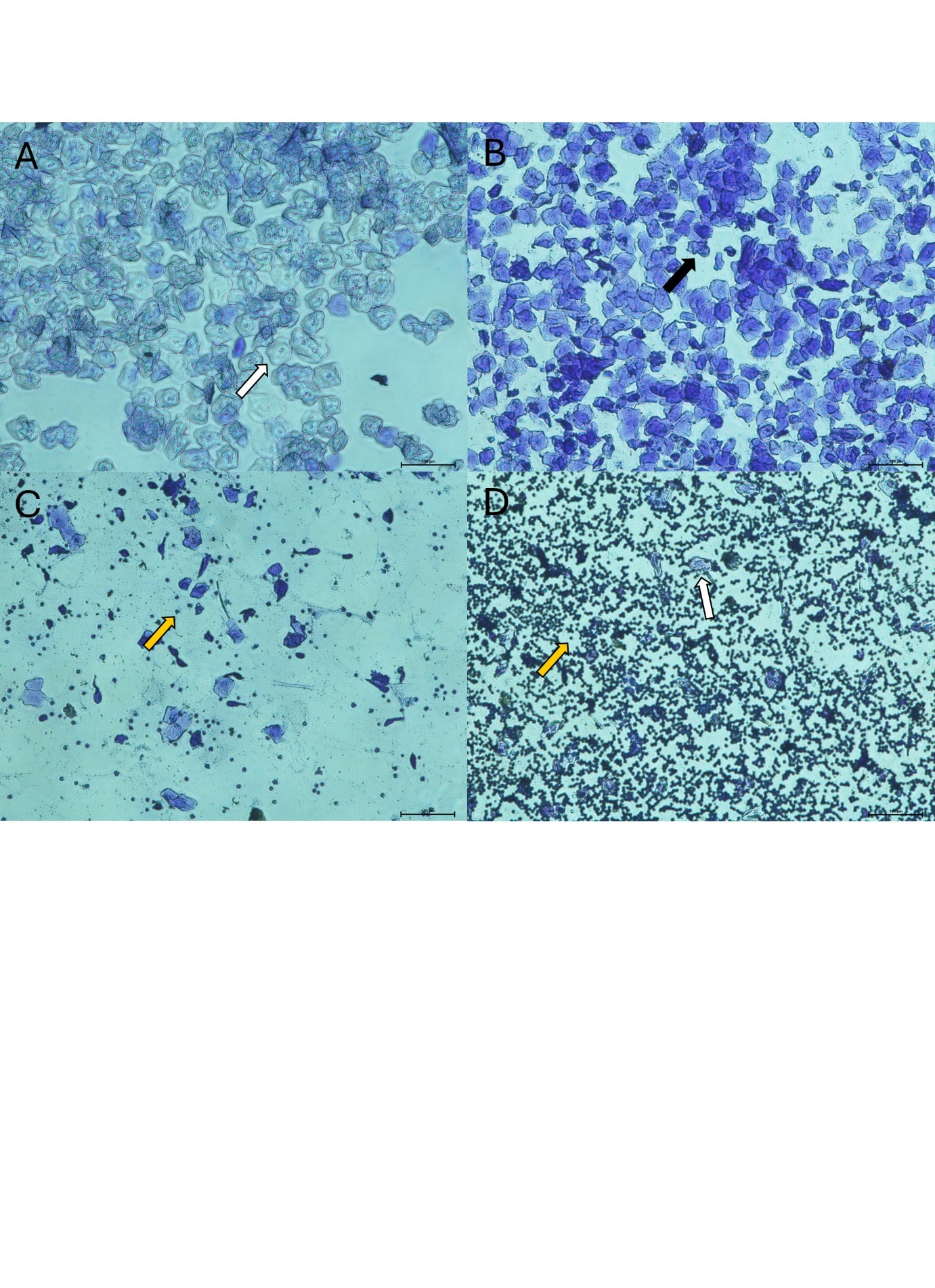

**Supplementary Figure 2. Estrus cycle phase determination from vaginal smears stained with crystal violet. (**A) The proestrus phase is characterized by the predominance of nucleated epithelial cells (white arrow). (B) The estrus phase is defined by the predominance of anucleated, cornified, irregularly shaped epithelial cells (black arrow). (C) Metestrus phase is identified by predominance of leukocytes (yellow arrow). (D) Diestrus phase is determined by the presence of both leukocytes (yellow arrow) and some nucleated epithelial cells (white arrow). Scale bar = 100 µm.
