## Supplementary Fig. S3 for "Maternal High-Fat Diet Induces Sex- and Estrous Cycle-Specific Glial Dysregulation in the Peripheral Offspring Retina"

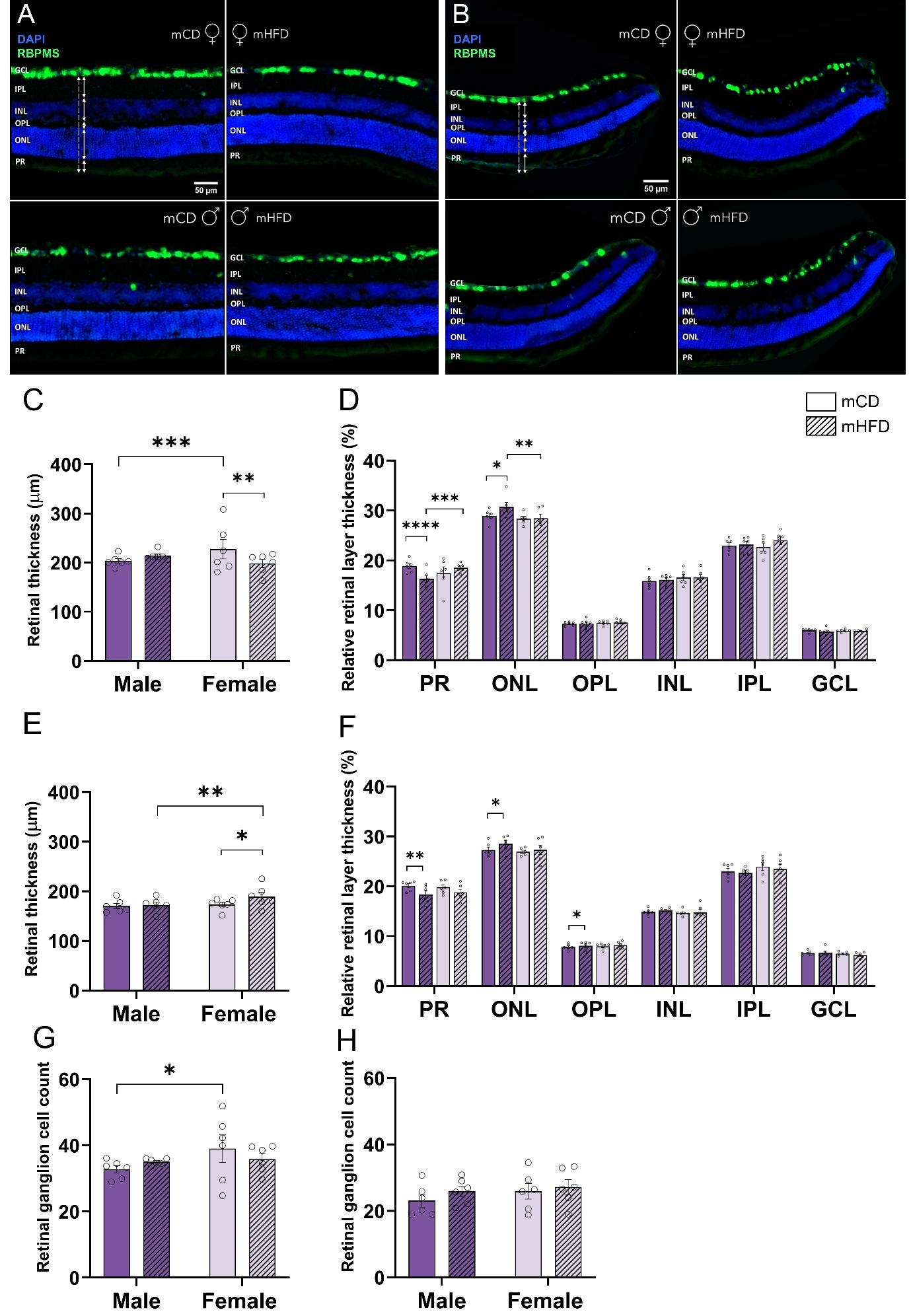

**Supplementary Figure 3. The effects of mHFD on the offspring’s retinal thickness and ganglion cell density.** Representative confocal images of RBPMS-labeled (green) retinal sections of central (A) and peripheral (B) retina from female (♀) and male (♂) offspring of dams who were fed either control (mCD) or high-fat diet (mHFD). GCL – ganglion cell layer. IPL – inner plexiform layer. INL – inner nuclear layer. OPL – outer plexiform layer. ONL – outer nuclear layer. PR – photoreceptor layer. Scale bar = 50 µm. (C, D) Retinal thickness (µm) and relative retinal layer thickness (%) in the central retina of the offspring. (E, F) Retinal thickness (µm) and relative retinal layer thickness (%) in the peripheral retina of the offspring. (G, H) Retinal ganglion cell number per 500 µm in the central and peripheral retina of the offspring. Mean ± SEM, n = 6 animals per group. *P*-values were assessed using nested two-way ANOVA with Tukey *post hoc* tests. **P* < 0.05, ***P* < 0.01, ****P* < 0.001, *****P* < 0.0001.
